# Characterization and Sap Transmission of Citrus Bent Leaf Viroid in Malaysia

**DOI:** 10.1101/751560

**Authors:** Yasir Iftikhar, Ying Wei Khoo, Taneswari Murugan, Nur Athirah Roslin, Rabiatul Adawiyah, Lih Ling Kong, Ganesan Vadamalai

**Author notes:** corresponding author: G. Vadamalai. All the authors contributed this work equally.

## Abstract

A 328 nucleotide (nt) variant of *Citrus bent leaf viroid* (CBLVd) was characterized from citrus varieties in Malaysia showing leaf bending, stunting and midvein necrosis. CBLVd was detected by RT-PCR assay using CBLVd specific primers in 12 out of 90 samples, collected from six different areas in Malaysia. The viroid was present in five species of citrus namely *Citrofortunella microcarpa, Citrus aurantifolia, C. hystrix, C. maxima* and *C. sinensis*. Sequence analysis of the isolates obtained from this study showed 99-100% sequence identity to CBLVd Jp isolate (AB006734). Inoculation of sap obtained from a CBLVd positive *C. aurantifolia*, inoculated into six months old *C. microcarpa* seedlings showed the symptoms leaf bending, reduced leaf size of matured leaves and mild mosaic between 4 to 6 months after inoculation. The presence of CBLVd in the inoculated seedlings were confirmed by RT-PCR assay and sequencing.

**Author Summary:** The authors during a limited survey collected the citrus samples from citrus growing areas of Malaysia to detect the citrus viroids. Citrus viroids are associated with decline in citrus production. Thus, the presence of the citrus viroids and their spread needs to be investigated to facilitate the management of this pathogen in citrus orchards. The authors detected and characterized Citrus bent leaf viroid in Malaysian citrus.

## Introduction

Citrus is an important economic fruit crop in Malaysia producing about 40,014 metric tons of citrus in 2016 (FAOSTAT, 2016). Citrus species such as *C. aurantifolia, C. hystrix, C. limon, C. maxima, C. microcarpa, C. reticulata* and *C. sinensis* are planted in Malaysia (Chooi, 1994; Md Othman et al. 2016). Citrus production is affected by various pest and diseases including viroid diseases. Citrus viroids cause devastating impact in citrus industries by reducing yield and plant health. To date, there are seven viroids from four genera of *Pospiviroidae* have been reported infecting citrus species. Major citrus viroids are Citrus exocortis viroid (CEVd) of *Pospiviroid*, Citrus bent leaf viroid (CBLVd), Citrus dwarfing viroid (CVd-III), Citrus viroid V and Citrus viroid VI (CVd-V and CVd-VI) of *Apscaviroid, Hop stunt viroid* (HSVd) of *Hostuviroid*, and Citrus bark cracking viroid (CBCVd) of *Cocadviroid* (Reanwarakon et al. 2003). Citrus viroids induce characteristic symptoms like dwarfing, leaf bending, mid vein and petiole necrosis (Ito et al 2001). In addition, some of other symptoms that have been associated with citrus viroids include wood pitting, bark scaling, leaf epinasty and stunting (Cao et al. 2009, 2010; Ito et al. 2001; Malfitano et al. 2005). The viroid infection in the citrus plants has been characterized as solely or in combination with each other (Hashemian et al. 2010). Among them, CEVd and CBLVd are widely distributed (Ito et al 2002).

CBLVd have been reported in Al-Ain, United Arab Emirates (UAE) (Al-Shariqi et al. 2013), Campania, South Italy (Malfitano et al. 2005), Kohgiluyeh-Boyerahmad, Iran (Mazhar et al. 2014) and Japan (Ito et al. 2000). It has different isolates which includes CVd-Ia, CVd-Ib and a distinct variant CVd-1-LSS (Ito et al. 2000). CVd-Ib (315-319 nt) was first sequenced and renamed as CBLVd, while CVd-Ia comprised of 327-329 nt. The CVd-I-LSS (325-330 nt) isolates have only 82-85% similarity in sequence with CVd-I variants and regarded as low sequence similarity. CBLVd and its isolates have been reported in Asia but but there were no reports of citrus viroids in Malaysia until recently, where CBLVd variants was been reported in Malaysian citrus (Khoo et al., 2017). This paper describes molecular detection and characterization of CBLVd from citrus species in Malaysia and its sap transmissibility to the healthy citrus plants.

## Results

### Detection of CBLVd using RT-PCR

RT-PCR amplification of total nucleic acid extracted from 133 symptomatic and 60 non-symptomatic citrus leaf samples using CBLVd specific primers showed the presence of CBLVd in 23 out of 133 symptomatic citrus samples collected from five states in Malaysia; four samples were from Johor, 10 from Malacca and nine from Selangor. All non-symptomatic citrus samples were not detected with CBLVd. Among the seven citrus species sampled, the CBLVd was detected six species including three *C. aurantiafolia* samples, two from *C. hystrix*, one from *C. jambhiri*, four from *C. maxima*, 11 from *C. microcarpa*, and two from *C. sinensis* (Table 1). CBLVd was not detected in *C. reticulata*. Positive samples produced a band approximately 330 bp in the agarose gel analysis (Fig. 1).

**Fig. 1.**
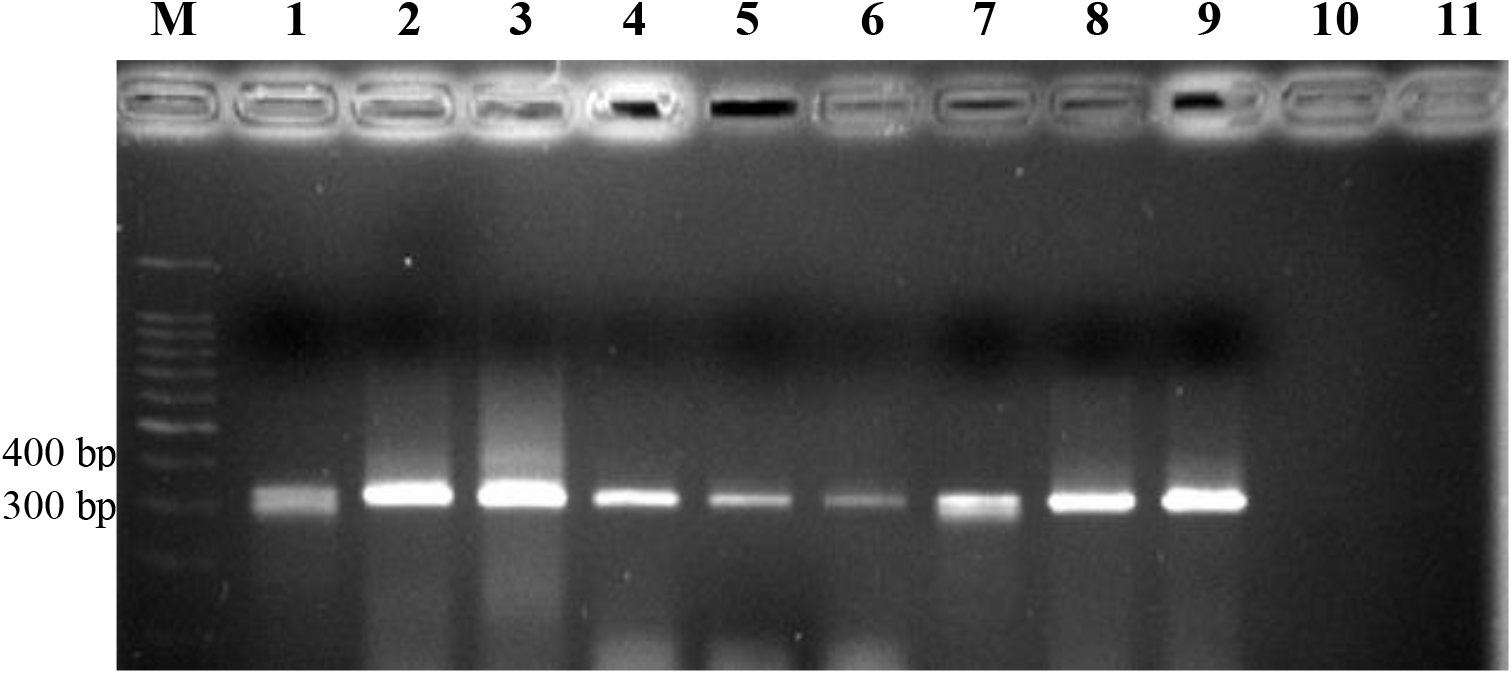
Reverse transcription-polymerase chain reaction assay of CBLVd positive samples from Johor, Malacca and Selangor with CBLVd specific primer sets analyzed in 2.0% agarose gel electrophoresis. An amplicon of approximately 330 bp were observed in CBLVd positive samples in Lane 1-3 (Johor), Lane 4-6 (Malacca), Lane 7-9 (Selangor). Lane 10 is non-template control. Lane 11 is healthy sample from non-symptomatic citrus leaf sample.

**Table 1.** Detection of infection of CBLVd in citrus species from commercial orchards in different states using Multiplex RT-PCR.

| State (N)<sup>a</sup> | Number of citrus species detected with CBLVd (n)<sup>b</sup> |  |  |  |  |  |  | Total |
| --- | --- | --- | --- | --- | --- | --- | --- | --- |
|  | *C. aurantifolia* | *C. hystrix* | *C. jambhiri* | *C. maxima* | *C. microcarpa* | *C. reticulata* | *C. sinensis* |  |
| **Johor (23)** | 2 (15) | 0 (4) | - | - | 0 (1) |  | 2 (3) | 4 |
| **Malacca (25)** | 1 (2) | - | 1 (1) | - | 8 (22) | - | - | 10 |
| **Selangor (48)** | 0 (8) | 2 (12) | - | 4 (7) | 3 (20) | 0 (1) | - | 9 |
| **Pahang (12)** | 0 (2) | 0 (7) | - | - | 0 (2) | - | 0 (1) | 0 |
| **Perak (25)** | - | - | - | 0 (8) | 0 (2) | 0 (15) | - | 0 |
| **Total** | 3 (27) | 2 (23) | 1 (1) | 4 (15) | 11 (47) | 0 (16) | 2 (4) | 23 |

“ - ”: absence of that particular citrus species in the field a in parentheses: total number of sampled plants from different states b in parentheses: number of sampled plants from each citrus species

#### Cloning and Sequencing

Cloning and sequencing of the purified PCR products of the RT-PCR positive samples produced 12 cDNA clones of 328nt. Sequence analysis of the 12 clones showed that all clones had 99-100% similarity with CBLVd isolate Japan (GenBank Acc No. AB006734). Five of the clones were 100% similar with CBLVd isolate Japan while seven clones were 99 % similar with the Japan isolate. Among the 12 clones, four clones were from Malacca (MyMalacca), four clones from Johor (MyMuar) and four clones from Selangor (MySerdang) (Table 2). In addition, out of the 12 clones, five were obtained from *C. microcarpa*, three clones from *C. aurantifolia*, and two clones from *C. sinensis* and *C. maxima* respectively (Table 2). All the sequences obtained in this study were deposited in GenBank (Table 2).

**Table 2.** Host, GenBank accession numbers and sequence similarity of CBLVd clones obtained in Malaysia.

| Clone | Host | GenBank<br>Accession<br>number | % identity with<br>CBLVd Japan<br>isolate<br>(AB006734) |
| --- | --- | --- | --- |
| MyMalacca01/08 | *Citrofortunella*<br>*microcarpa* | MF361856 | 100 |
| MyMalacca01/10 | *Citrofortunella*<br>*microcarpa* | MF346702 | 99 |
| MyMalacca01/13 | *C. aurantifolia* | MF361857 | 100 |
| MyMalacca01/20 | *Citrofortunella*<br>*microcarpa* | MF346703 | 99 |
| MyMuar01/01 | *C. aurantifolia* | KX823338 | 100 |
| MyMuar01/03 | *C. aurantifolia* | KX823339 | 99 |
| MyMuar01/14 | *C. sinensis* | KU194472 | 100 |
| MyMuar01/15 | *C. sinensis* | KU936033 | 99 |
| MySerdang01/01 | *C. maxima* | KX823340 | 99 |
| MySerdang01/02 | *C. maxima* | KX823341 | 99 |
| MySerdang01/06 | *Citrofortunella*<br>*microcarpa* | KX823342 | 99 |
| MySerdang01/08 | *Citrofortunella*<br>*microcarpa* | KX823343 | 100 |

Alignment of the seven clones with 99% sequence similarity showed base substitutions in Terminal left (TL), Pathogenic (P), Variable (V) and Terminal right (TR) domain of the CBLVd Jp isolate (Table 3). Clone MySerdang01/01 had a substitution at base 59 (A→G) of the P region. Meanwhile, clone MySerdang01/02 had a substitution at base 33 (C→U) and 251 (U→C), 265 (U→C) of TL and P region respectively. Base substitutions also occurred in clone MySerdang01/06 at 285 (A→G) and 177 (C→U) at P and TR region respectively. The base substitution that occurred in MyMuar01/03 was at base 69 (G→C) and 208 (C→A), 209 (G→A) of the P and V region respectively. Meanwhile, the base substitution of clone MyMuar01/15 was similar to clone MyMuar01/03, but with an addition of a substitution at base 118 (G→A) of V region. Base substitution also occurred in clone MyMalacca01/10 at base 69 (G→C) of P region while for clone MyMalacca01/20, the base substitution occurred at 180 (A→G) of TR region (Table 3).

**Table 3.** Nucleotide changes of CBLVd clones obtained in this study compared with CBLVd isolate Japan (AB006734)

| Isolate | Position of nucleotides changes in compared with<br>CBLVd isolate Japan (GenBank Acc no AB006734) |  |  |  |
| --- | --- | --- | --- | --- |
|  | TL | P | V | TR |
| MySerdang01/01<br>(KX823340) | - | 59 (A→G) | - | - |
| MySerdang01/02<br>(KX823341) | 33 (C→U) | 251 (U→C),<br>265 (U→C) | - | - |
| MySerdang01/06<br>(KX823342) | - | 285 (A→G) | - | 177 (C→U) |
| MyMuar01/03<br>(KX823339) | - | 69 (G→C) | 208 (C→A),<br>209 (G→A) | - |
| MyMuar01/15<br>(KU936033) | - | 69 (G→C) | 118 (G→A),<br>208 (C→A),<br>209 (G→A) | - |
| MyMalacca01/10<br>(MF346702) | - | 69 (G→C) | - | - |
| MyMalacca01/20<br>(MF346703) | - | - | - | 180 (A→G) |

#### Phylogenetic analysis

Phylogenetic tree demonstrated the sequences were grouped into four main clades that exhibited the origin of samples. All Malaysian clones were in clade I. MyMalacca01/08, MyMalacca01/13, MyMuar01/01, MyMuar01/14, MySerdang01/08, and CBLVd Japan isolate (AB006734) were in sub-cluster A. MyMalacca01/10 was in sub-cluster B. MyMuar01/03 and MyMuar01/15 shared the same sub-cluster C. Sub-cluster D contained MyMalacca01/20, MySerdang01/02, MySerdang01/06 and MySerdang01/01 (Fig. 2).

**Fig. 2.**
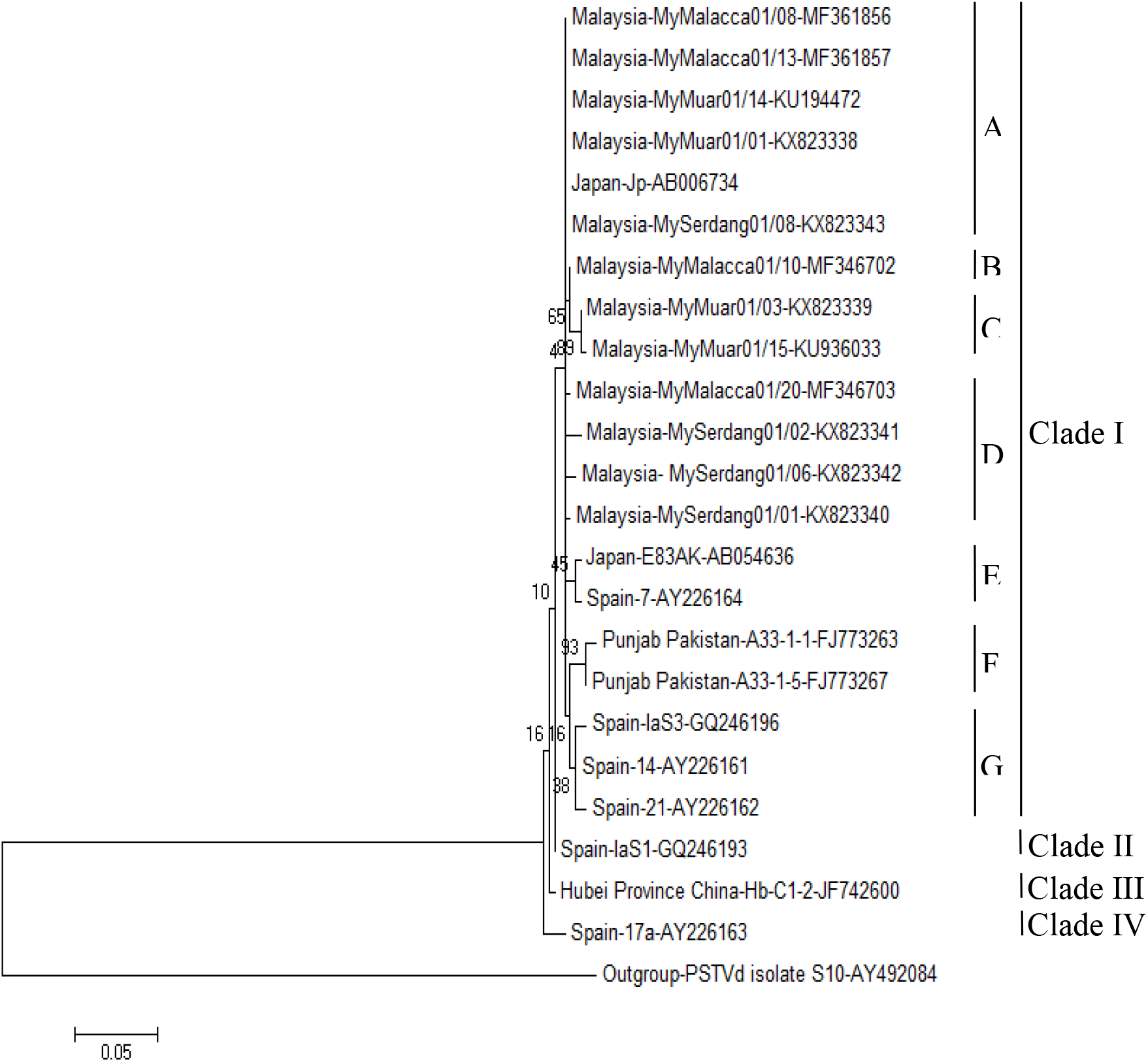
Phylogenetic relationships of Citrus bent leaf viroid (CBLVd) sequence variants using MEGA 6.0 based on 1000 bootstrap replication. Potato spindle tuber viroid (PSTVd, Accession No. AY492084) was used as outgroup sequence.

#### Pathogenicity test

All the 10 calamondin (*C. microcarpa*) seedlings inoculated with the sap extracted from a CBLVd positive citrus plant (MySerdang01/08) exhibited leaf bending, leaf rolling, chlorosis and mild necrosis on mid rib of the leaves. In addition, the matured leaves of inoculated seedlings were smaller in size compared to the healthy seedlings. However, no stunting was observed in all inoculated seedlings. All the symptoms were observed after three month post inoculation (Fig. 3). Leaf samples from all the 10 inoculated seedlings were detected with CBLVd through RT-PCR using CBLVd specific primers (Fig. 4). The presence of CBLVd was confirmed through sequencing the PCR product which resulted in 100% similarity with the sequence of CBLVd Japan isolate (AB006734).

**Fig. 3.**
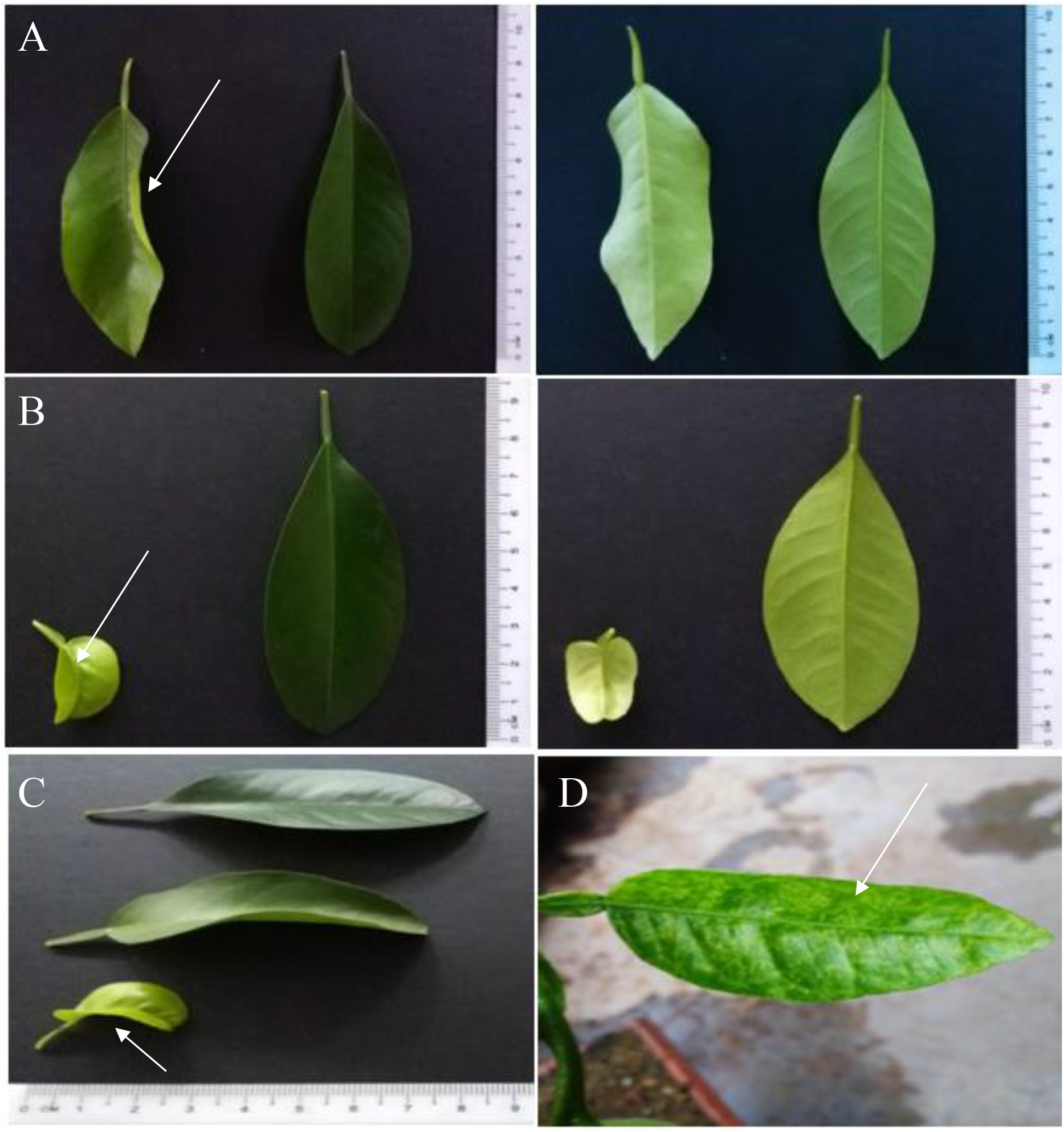
Symptoms observed on 6 month old calamondin seedlings were indicated with arrow. A, B, C and D, Leaf rolling (A), Small leaves (B), mild necrosis on mid rib (B), leaf bending (C), chlorosis (D) induced by CBLVd in 6-month-old calamondin seedlings after 3-month post-inoculation.

**Fig. 4.**
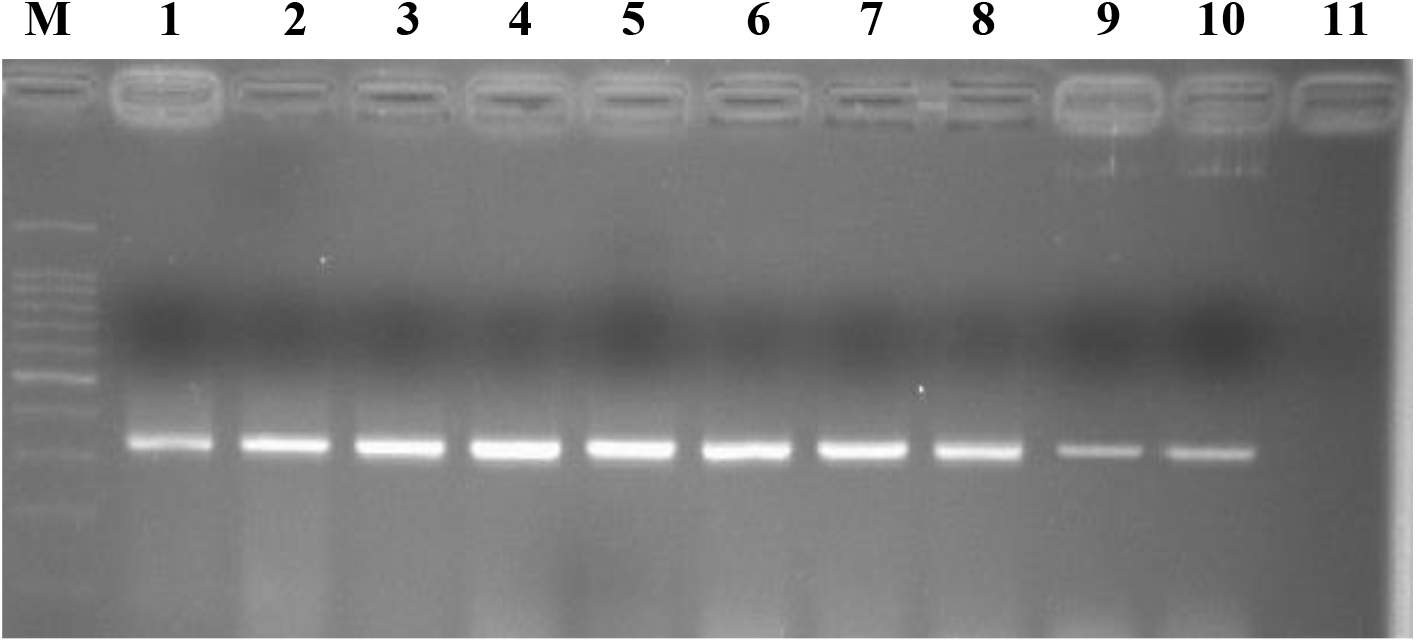
Reverse transcription-polymerase chain reaction assay of CBLVd positive samples from ten CBLVd sap-inoculated seedlings with CBLVd specific primer sets analyzed in 2.0% agarose gel electrophoresis. An amplicon of approximately 330 bp were observed in ten inoculated seedlings samples in Lane 1-10. Lane 11 is healthy distilled water inoculated seedlings sample.

## Discussion

RT-PCR amplification of total nucleic acid extracted from citrus leaf samples using CBLVd specific primers showed the presence of CBLVd in 23 samples collected from 3 states in Malaysia and six citrus species. However, as only 23 out of 133 symptomatic citrus plants were detected with CBLVd, which indicates that the symptoms observed in the field were not specific for CBLVd infection. The field symptoms observed might be due other causes as Citrus Tristeza Virus also cause stunting (Moreno *et al*., 2008) and leaf yellowing could be due to nitrogen deficiency in citrus plants (Futch and Tucker, 2003). CBLVd might be endemic in Asia Pacific region. Previous studies have shown that CBLVd was widely distributed in Asia Pacific region such as China, Japan, Malaysia, Pakistan and Philippines (Hataya et al. 1998; Ito et al. 2000; Cao et al. 2009; Wu et al. 2014; Khoo et al. 2017). Results showed that CBLVd were detected in *Citrofortunella microcarpa, C. aurantiafolia, C. hystrix, C. maxima* and *C. sinensis* but not in *C. reticulata*. The findings suggested that citrus could be a natural host to CBLVd (Eiras et al. 2013; Hammond and Owens, 2006). CBLVd was detected in Johor, Malacca and Selangor suggested that infected citrus plant materials could be the source of CBLVd dissemination. Previous studies have showed that contaminated propagated materials helped in the dissemination of Citrus greening and Citrus tristeza virus in Peninsular Malaysia (Ahmad et al. 2008; Ayazpour et al. 2011). However, further studies are required to determine the epidemiology of CBLVd in Peninsular Malaysia. Interestingly, CBLVd was not detected in citrus plants in Pahang and Perak. Exploration about the source of propagated materials and rootstock used for citrus plants in Perak and Pahang should be examined.

CBLVd of 328 nt was detected by RT-PCR using CBLVd specified primer sets indicated the efficiency of RT-PCR to detect and characterize CBLVd. Bernard and Durán-Vila (2006) have reported about RT-PCR being employed to detect and characterize citrus viroids. Sequence analysis of 12 clones showed that the CBLVd variants from this study showed 99-100% sequence homology to CBLVd isolate Japan (AB006734). Besides, the clones isolated from *C. microcarpa, C. aurantiafolia, C. maxima* and *C. sinensis* were identical and were defined as the consensus sequence of CVd-Ia which is a derivative of citrus bent leaf viroid (Formerly known as CVd-Ib). Therefore, based on the percentage sequence similarity, the CBLVd variant from citrus in Malaysia was determined as CVd-Ia (Hataya et al. 1998; Ito et al. 2002). In addition, sequence analysis showed that seven out of 12 clones obtained in this study had base substitution mainly occurred in Pathogenicity and Variable domain compared to CBLVd japan isolate. Substitution of nucleotide occurring at different domains on the secondary structure of the viroid, suggesting the adaptation of CBLVd to the host condition. The finding is consistent with Lin *et al*. (2015), which showed that nucleotides alterations were used by two citrus viroids (CEVd and HSVd) to adapt to different hosts.

The phylogenetic analysis revealed that the 12 CBLVd isolates obtained in this study shared the sub-cluster with isolates from Japan (AB006734) indicating the closeness of region shaped genetic homology. This finding is consistent with previous study revealed that some of Australian grapevine viroid (AGVd) isolates from India similar to Chinese isolates (Adkar-Purushothama et al. 2014). No information available with regards to the molecular phylogenetic analyses to adequately identify and illuminate the evolution of CBLVd from Japan with relation to the Malaysian isolates and other isolates from different countries. Extensive phylogenetic analyses are required to give more information on the source of origin as well as evolutionary relationship of CBLVd.

Inoculation of calamondin (*C. microcarpa*) seedlings with sap extracted from a calamondin (MySerdang01/08) induced symptoms such as leaf bending, leaf rolling, chlorosis and mild necrosis on mid rib of the leaves, 3-month post inoculation. The symptoms observed in this study, specifically the leaf bending symptoms on the inoculated calamondins corresponds to the symptoms induced by CBLVd infections in citrus regardless of varieties as previously reported (Gandía and Durán-Vila, 2004). In addition, the observation of chlorosis, mid vein necrosis, leaf rolling and smalling of matured leaves in this study have not been reported in calamondins. The appearance of these symptoms could be attributed to the nucleotide changes of CBLVd in host calamondin (Sano et al. 1992). This study also showed that CBLVd is able to be transmitted through sap. The mechanical transmission of viroids through vegetative propagation and contact of tools used in plantations has already been described by Hadidi et al. 2003, thus, the current finding could serve as a platform for future study of interlink between sap transmission and citrus viroids and potential insect vector.

In conclusion, CBLVd was detected from citrus in Malaysia using RT-PCR with CBLVd specific primers. CBLVd was detected in *C. aurantiafolia, C. hystrix, C. jambhiri, C. maxima, C. microcarpa*, and *C. sinensis* except *C. reticulata*. Cloning and sequencing showed that all CBLVd were variant of CVd-Ia with 328 nt in length with 99-100% sequence homology with CBLVd isolate Jp (AB006734). Pathogenicity of CBLVd in *C. microcarpa* was confirmed through mechanical inoculation of sap. Further studies on different variants of CBLVd, host range and geographical distribution are important to fill up the gap in research on CBLVd.

## Materials and Methods

### Collection of samples

Citrus samples were collected from citrus growing areas of Johor, Malacca, Pahang, Perak and Selangor. Leaves samples citrus plants exhibiting leaf bending, stunted growth and midvein necrosis were collected from 133 citrus plants of different varieties of citrus including calamondin (*Citrofortunella microcarpa* (Bunge) Wijnands), kaffir lime (*Citrus hystrix* DC.), key lime (*C. aurantifolia* (Cristm.) Swingle), mandarin (*C. reticulata* Blanco), pomelo (*C. maxima* Merr.), rough lemon (*C. jambhiri* Lush.) and sweet orange (*C. sinensis* (L.) Osbeck). In addition, 60 non-symptomatic citrus samples were also collected from the same citrus growing areas. The samples were surface sterilized with 10% sodium hypochlorite followed by distilled water and kept at −20°C until use.

### Extraction of total nucleic acid

The total nucleic acid was extracted from leaf samples using the TESLP buffer (0.13 M Tris-HCl (pH 8.9), 0.017 M EDTA (pH 7.0), 1.0 M LiCl, 0.83% SDS, 5% PVP) (Ito et al. 2000) following the steps described in Nakahara et al. (1999) with slight modification. Mid-ribs from 4 g of leaves were removed. The leaves were chopped into small pieces and grounded in liquid nitrogen using mortar and pestle. The powder was transferred to 50 ml screw cap tube. A total of 10 ml of TESLP buffer was added into the tube followed by 16 μl of 2-mercapthoethanol and incubated for 30 min at room temperature in a rotary shaker. The mixture was centrifuged at 13,000 *g* for 15 min. The resulting supernatant was collected and extracted with 2 vol of Phenol, Chloroform and Isoamyl Alcohol (PCA, 25:24:1). The mixture was again centrifuged for 15 min at 13,000 g. The supernatant was re-extracted into a new tube with 3 vol of Chloroform and Isoamyl Alcohol (CA, 24:1) followed by centrifugation for 15 min at 13,000 g. The supernatant was transferred into a new 15 ml screw cap tube added with 0.9 volume of 90% isopropanol. The mixture was mixed by inverting and incubated at −80°C for 30 min. The pellet was collected by centrifugation 15 min at 13,000 g. The pellet was washed twice with 1 ml of 70 % ethanol, air dried for 20-30 min and suspended in 50 μl of sterile double distilled water.

### Reverse transcription for RT-PCR

cDNA was synthesized using a reverse primer, CBLV-CP (5’-CGTCGACGAAGGCTCGTCAGCT-3’) (Ito et al. 2002) to synthesize the cDNA. Total nucleic acid (5.0 μl), reverse primer (1.0 μl) and double distilled water (2.5 μl) was added to a reaction vol of 8.5 μl. The reaction mix was incubated at 80°C for 12 min and immediately transferred to ice for 5 min. AMV-RT (1.0 μl), dNTPs (2.0 μl), RNAse Inhibitor (0.5 μl), MgCL_2_ (4.0 μl) and RT buffer (4.0 μl) were then added. The reaction was incubated at 55°C for 30 min and stopped by incubation at 10°C. The cDNA obtained was stored in −80°C freezer until use.

### Reverse Transcription Polymerase Chain Reaction Amplification

A set of specific primer to amplify the full-length genome of CBLVd, forward, CBLV-CM (5’-ACGACCAGTCAGCTCCTCTG-3’) and reverse, CBLV-CP (5’-CGTCGACGAAGGCTCGTCAGCT-3’) was used for RT-PCR (Ito et al. 2002). The final volume of PCR product was 25.0 μl which consisted of 12.5 μl of PCR master mix, 5.0 μl of cDNA, 5.5 μl of sterile double distilled water, 1.0 μl each for forward and reverse primers (0.4 pmol/ μl each primer in final concentration). The conditions for PCR amplification for 35 cycles were 94°C for 10 min, 94°C for 30 sec followed by gradual decreasing at 60°C for 1 min. Annealing was at 60°C for 10 sec. The reaction was ended with the extension at 72°C for 5 min.

### Agarose Gel Electrophoresis

The PCR products were fractionated in 2% agarose gel prepared in 1x TBE buffer at 100 V for 50 min. The gel was stained with ethidium bromide for 10 min followed by destained with distilled water for 5 min. The gel was visualized under Trans UV and captured with Gel Doc XR model.

### Cloning and sequencing

Positive PCR products were purified using MinElute^®^ Gel Extraction Kit (QIAGEN) according to the manufacturer’s recommendation. The purified PCR products were ligated into a cloning vector (pCR2.1-TOPO, Invitrogen) followed by transformation with heat shock at 42°c for 30 sec in a water bath. The samples were immediately put on ice for 5 min. SOC medium was added and incubated at 37°C for 90 min at 200 rpm. X-gal was spread gently on LBA plate containing 50 mg/ L kanamycin. The plate was sealed, wrapped with aluminum foil, inverted and incubated for 45 min prior to spreading the samples. The clones were selected and sequenced at First Base and Next Gene (Malaysia). The sequences were compared for similarity against the available sequences from NCBI database Blast program. The sequences from clones were aligned using the software BioEdit version 7.2.5 (Hall et al. 1999).

### Phylogenetic analysis

A phylogenetic tree was constructed by MEGA 6.0 employing the maximum likelihood method with bootstrap analysis (1,000 replicates) (Lin et al. 2015). Potato spindle tuber viroid (PSTVd, Accession No. AY492084) was used as outgroup sequence. The Malaysian isolates were analysed together with isolates from China, Japan, Pakistan and Spain (Table 4).

**Table 4.** Host, origin and sequence similarity of clones of CBLVd worldwide isolates with CBLVd isolate Japan (AB006734) from *C. clementina*.

| Clone | Host | Origin | Accession numbers | % identity with Jp isolate |
| --- | --- | --- | --- | --- |
| E83AK | ‘Satsuma’ mandarin orange | Japan | AB054636 | 99 |
| Jp | ‘Clementine de Nules’ mandarin orange | Japan | AB006734 | 100 |
| 7 | Citron | Spain | AY226164 | 99 |
| 14 | Citron | Spain | AY226161 | 99 |
| 17a | Citron | Spain | AY226163 | 98 |
| 21 | Citron | Spain | AY226162 | 99 |
| IaS1 | Clementine mandarin | Spain | GQ246193 | 99 |
| IaS3 | Clementine mandarin | Spain | GQ246196 | 99 |
| A33-1-1 | Sweet orange | Punjab, Pakistan | FJ773263 | 98 |
| A33-1-5 | Sweet orange | Punjab, Pakistan | FJ773267 | 98 |
| Hb-C1-2 | Citron | Hubei Province, China | JF742600 | 99 |

### Pathogenicity Test

Leaf sap was extracted from 5 g of CBLVd positive citrus leaves (MySerdang01/08) exhibiting leaf bending, petiole necrosis and yellowing symptoms in 0.02 M phosphate buffer (pH 7) with one vol of tissue to five vol of buffer using pestle and mortar (Iftikhar et al. 2004). The sap was filtered through muslin cloth and kept on ice. Carborandum was dusted on 6 months old calamondin (*C. microcarpa*) seedlings before inoculation. Sap was applied to the leaves. About 4-6 leaves of each seedling were selected for inoculation. Altogether 10 seedlings were inoculated with filtered sap. The inoculated seedlings were washed out under the tap water after 3-5 min to remove the excessive filtrate. Another ten seedlings were inoculated with distilled water as control. All seedlings were kept in insect free screenhouse for 6 months to observe for symptom development and CBLVd presence.

## Acknowledgements

The authors gratefully acknowledge the collaboration of TWAS and UPM under TWAS-UPM post-doc fellowship program and University of Sargodha, Pakistan for sabbatical leave to conduct this study.

